# A honey bee symbiont buffers larvae against nutritional stress through lysine supplementation

**DOI:** 10.1101/2022.01.11.475899

**Authors:** Audrey J. Parish, Danny W. Rice, Vicki M. Tanquary, Jason M. Tennessen, Irene L.G. Newton

## Abstract

Honey bees, the world’s most significant agricultural pollinator, have suffered dramatic losses in the last few decades (1,2). These losses are largely due to the synergistic effects of multiple stressors, the most pervasive of which is limited nutrition (3–5). The effects of poor nutrition are most damaging in the developing larvae of honey bees, who mature into workers unable to meet the needs of their colony (6–8). It is therefore essential that we better understand the nutritional landscape experienced by honey bee larvae. In this study, we characterize the metabolic capabilities of a honey bee larvae-associated bacterium, *Bombella apis* (formerly *Parasaccharibacter apium),* and its effects on the nutritional resilience of larvae. We found that *B. apis* is the only bacterium associated with larvae that can withstand the antimicrobial larval diet. Further, we found that *B. apis* can synthesize all essential amino acids and significantly alters the amino acid content of synthetic larval diet, largely by increasing the essential amino acid lysine. Analyses of gene gain/loss across the phylogeny suggest that two distinct cationic amino acid transporters were gained by *B. apis* ancestors, and the transporter LysE is conserved across all sequenced strains of *B. apis.* This result suggests that amino acid export is a key feature conserved within the *Bombella* clade. Finally, we tested the impact of *B. apis* on developing honey bee larvae subjected to nutritional stress and found that larvae supplemented with *B. apis* are bolstered against mass reduction despite limited nutrition. Together, these data suggest an important role of *B. apis* as a nutritional mutualist of honey bee larvae.

## Introduction

All metazoan evolution has occurred in the context of microorganisms, which exist in and around all living things. Consequently, mutualism between microbes and animal hosts is an ancient and widespread phenomenon (see (9)). Bacterial symbionts can have dramatic effects on animal hosts, including nutrient supplementation of incomplete host diets (10–13), protection from parasites and pathogens (14–16), and providing developmental cues (14, 17–21). Many eukaryotic hosts rely on bacterial partners for fundamental aspects of their metabolism, such as providing key nutrients absent or insufficient in the host diet (12, 22–24). In fact, the ecological variety of insects on earth is due in part to their ability to form new niches in previously inhospitable nutritional conditions, an accomplishment often achieved via association with bacterial partners (25–27). The European honey bee, *Apis mellifera,* is an excellent exemplar of this phenomenon, whereby the gut microbiome allows the colony to subsist upon recalcitrant plant pollen and nectar (28–31).

Honey bees are essential for pollinating food crops, resulting in a multi-billion-dollar global industry(32). However, managed honey bee colonies have suffered substantial losses in the last two decades (1). US beekeepers reported losing 40.5% of their managed colonies between 2015 and 2016 alone (2). These declines are often credited to a combination of stressors, the crux of which is poor nutrition (3–5). Poor nutrition for managed honey bee colonies is partly due to dwindling natural floral resources and an increase in reliance on large monoculture crops (33, 34). These large monocultures pose especially difficult nutritional landscapes for honey bees, as pollen from most individual crops only barely provides a colony’s base nutritional requirements (35, 36). This discrepancy between available and required protein is most problematic for larvae, the immature stage of honey bee workers (6). Ample multifloral pollen protein is required for honey bee larval development, and insufficient protein during this stage can have cascading effects through the colony (6, 7). Larvae deprived of adequate protein mature into adults who are stunted in size and deficient in their ability to forage for floral resources, further exacerbating nutritional stress to the next generation of brood and compromising colony dynamics (7, 8). Additionally, honey bees raised on pollen containing inadequate protein are significantly more likely to fall victim to secondary stressors such as viral pathogens, *Varroa destructor* mites, and *Nosema apis* infection (4, 5, 37, 38).

However, the microbiome can significantly modulate larval nutrition in holometabolous insects (12, 39, 40). The adaptive decoupling of larval and adult phases, which dedicates the larval phase to growth, creates an opportunity for growth-promoting nutritional symbionts to associate specifically with larvae (41,42). The most extensively studied examples of nutritional symbionts of larvae come from studies in *Drosophila melanogaster,* where just a single member of the microbiome can rescue larval growth despite severe protein limitation (17, 43). Yet less is known about how the microbial communities associated with honey bee larvae contribute to their nutrition and development. Honey bee larvae are nurtured by their adult nestmates, who feed them a larval diet of nectar, pollen, and royal jelly (44). This larval diet is strikingly low in bacterial diversity and is occupied predominantly by the bacterium *Bombella apis* (formerly *Parasaccharibacter apium), Lactobacillus,* and *Fructobacillus* species (45, 46). *B. apis* is consistently associated with honey bee larvae, larval diet, and the adult glands which secrete royal jelly, but is not found in large numbers in the adult worker gut (45–48). Therefore, *B. apis* seems particularly well positioned to serve a nutritional role in honey bee larval development.

Here, we present data showing that indeed, *B. apis* is supplementing honey bee larvae through its secretion of essential amino acids. We first asked which bacterial members of the honey bee larval microbiome community can survive in the *in vitro* larval diet by subjecting a panel of strains to media containing a gradient of royal jelly. We subsequently performed a comparative genomic analysis across all sequenced strains of *Bombella* and related *Saccharibacter* to identify significant gene conservation and gene gain/loss. This study confirmed that all *B. apis* strains can synthesize all amino acids and have acquired multiple amino acid transporters. We then selected one strain of *Bombella apis,* A29, and performed a microbiological and metabolomic analysis of its metabolic potential as a nutritional mutualist. Finally, we modified an established *in vitro* larval rearing protocol to measure the effect of *B. apis* dietary supplementation on the growth of larvae experiencing nutritional stress. Our results strongly support the hypothesis that *B. apis* is a nutritional symbiont of honey bee larvae.

## Results and Discussion

### Only *Bombella apis* persists in honey bee larval diet

The honey bee larval diet comprises nectar, pollen, and royal jelly (6, 49). Royal jelly has long been known to possess potent antimicrobial properties, due to its acidity, viscosity, and the presence of antimicrobial peptides (50–52). Therefore, any bacterial mutualist exposed to the larval diet must be capable of tolerating this strongly antimicrobial environment. The honey bee larval microbiome has been characterized using 16S rRNA gene amplicons and comprises a limited number of bacterial taxa, predominately *Bombella apis, Lactobacillus kunkeei,* and *Fructobacillus fructosus* (plus uncharacterized *Lactobacillus* spp and *Fructobacillus* spp, and occasionally *Bifidobacterium* spp) (47, 53). To assess the ability of larvae-associated microbes to survive in royal jelly, we subjected bacterial strains to media containing a gradient of royal jelly (up to 50%) for 24 hours and determined strain persistence by counting resulting CFUs (Figure 1A). The 50% royal jelly treatment recapitulates the highest proportion of royal jelly used in standard *in vitro* larval rearing diets (54, 55). The bacterial media used here and in following assays was developed by our lab for the rapid growth of larvae-associated strains and is based on the components of the honey bee larval diet (Bacto-Schmehl (BS), see Methods). All five strains of *Bombella apis* assayed were able to survive at all levels of royal jelly (Figure 1B). Some *B. apis* strains show a dip in the number of CFUs recovered between the media control and lowest concentration of royal jelly, indicating a degree of susceptibility to royal jelly inhibition (Figure 1C). CFUs varied between strains, with B8 and C6 displaying the least reduction when royal jelly is first introduced. A29 and MRM1T show the greatest sensitivity to royal jelly addition, each exhibiting a 10-fold reduction in CFUs compared to media alone. In contrast, *Lactobacillus kunkeei* AJP1 and *Fructobacillus fructosus* AJP3 were highly sensitive to royal jelly addition. *L. kunkeei* AJP1 displayed a dose-dependent decline in the number of CFUs recovered at each increase in royal jelly in the media, with near total inhibition at 50%. *F. fructosus* AJP3 was unable to survive in any concentration of royal jelly beyond 10%. Overall, *B. apis* survival when challenged with royal jelly appears robust compared to other bacteria often identified as larvae-associated.

**Figure 1.**
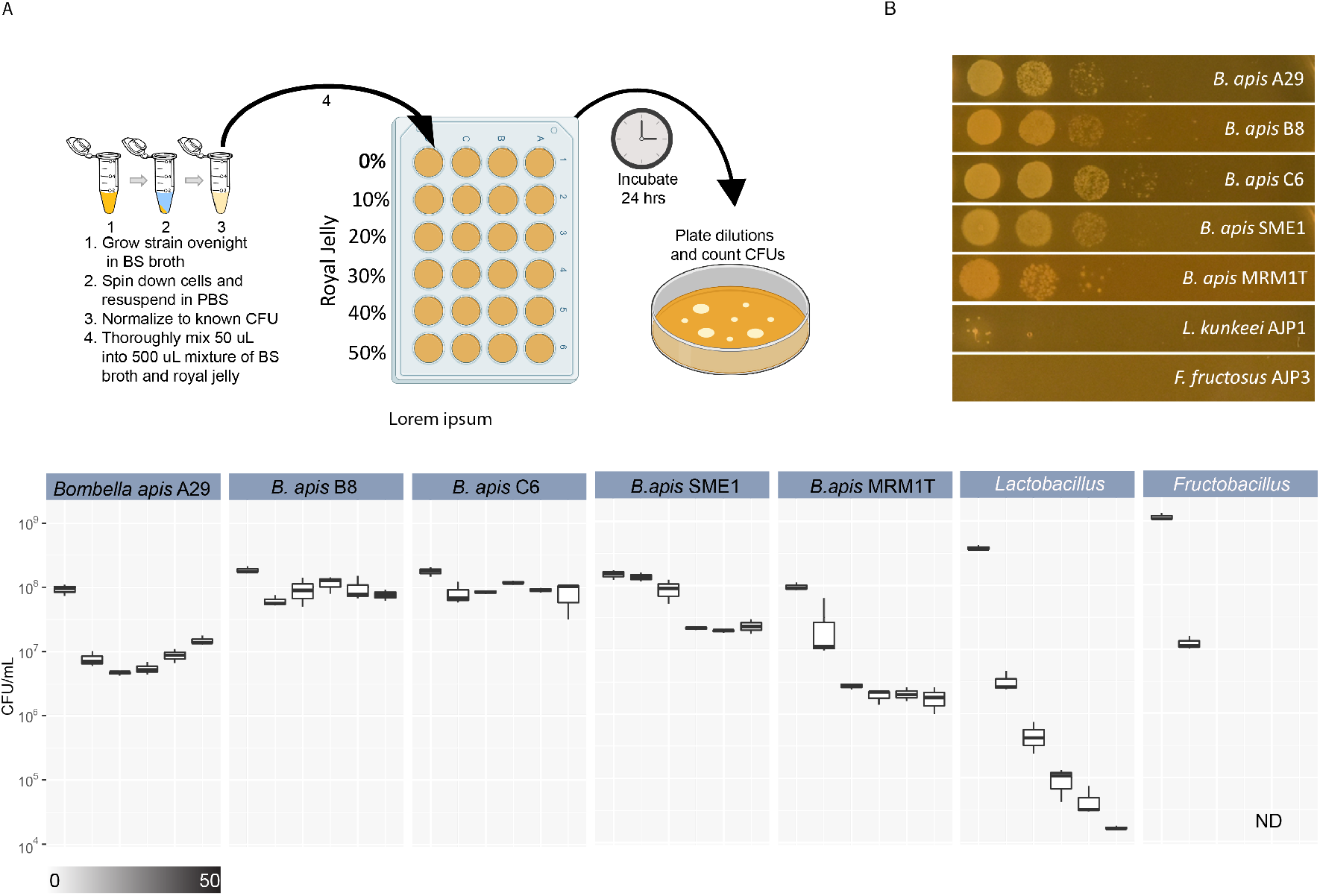
Only *Bombella apis* can tolerate the antimicrobial properties of royal jelly. **A.** The survival of a panel of larvae-associated bacteria in the presence of royal jelly was assessed by subjecting each to media containing a gradient of royal jelly from 0-50%. Strains were incubated overnight and plated on agar media to count CFUs. **B.** Representative images of the spot-dilution plates used to count CFUs after incubation in 50% royal jelly. **C.** Boxplots containing the total counts of CFUs resulting from each strain across all concentrations of royal jelly. Each concentration of all strains was calculated across three biological replicates.

All bacteria assayed here are associated with the larval diet niche from previous sequencing studies, but we do not know if the DNA sequenced arose from living cells or was ephemerally maintained from cells lysed in the diet (47, 53). Our findings suggest that *Bombella apis* and, to a lesser degree, *Lactobacillus kunkeei* are living in this niche, and other frequently sequenced bacteria such as *Fructobacillus fructosus* are likely not. This is in line with previous genomic evidence establishing *B. apis* as a honey bee-associated bacterium (56). Hosts who associate with horizontally acquired symbionts require a means of winnowing beneficial partners from environmental microbes, a phenomenon that has been predominantly studied in the context of selectivity of host tissues (57–60). However, honey bees mature in a built environment which can itself exert selective pressures on bacterial assemblages. In the case of the larval niche, it appears that the presence of royal jelly selects strongly for *Bombella apis* strains to associate with growing larvae.

### *B. apis* can produce all essential amino acids

As *B. apis* can persist in larval diet, we reasoned that it may be able to metabolically transform it. To first consider how *B. apis* may modify the host diet, we used tools provided by the DOE’s (IMG/M) website to explore the metabolic capabilities of a sequenced strain, A29. The *B. apis* genomes encode the complete biosynthetic pathways required to produce all canonically essential amino acids (Figure 2A). We then extended this result to all sequenced *B. apis* strains using an analysis of conserved orthologs; all sequenced *B. apis* strains retain the ability to synthesize all amino acids (Supplementary Table 1).

**Figure 2.**
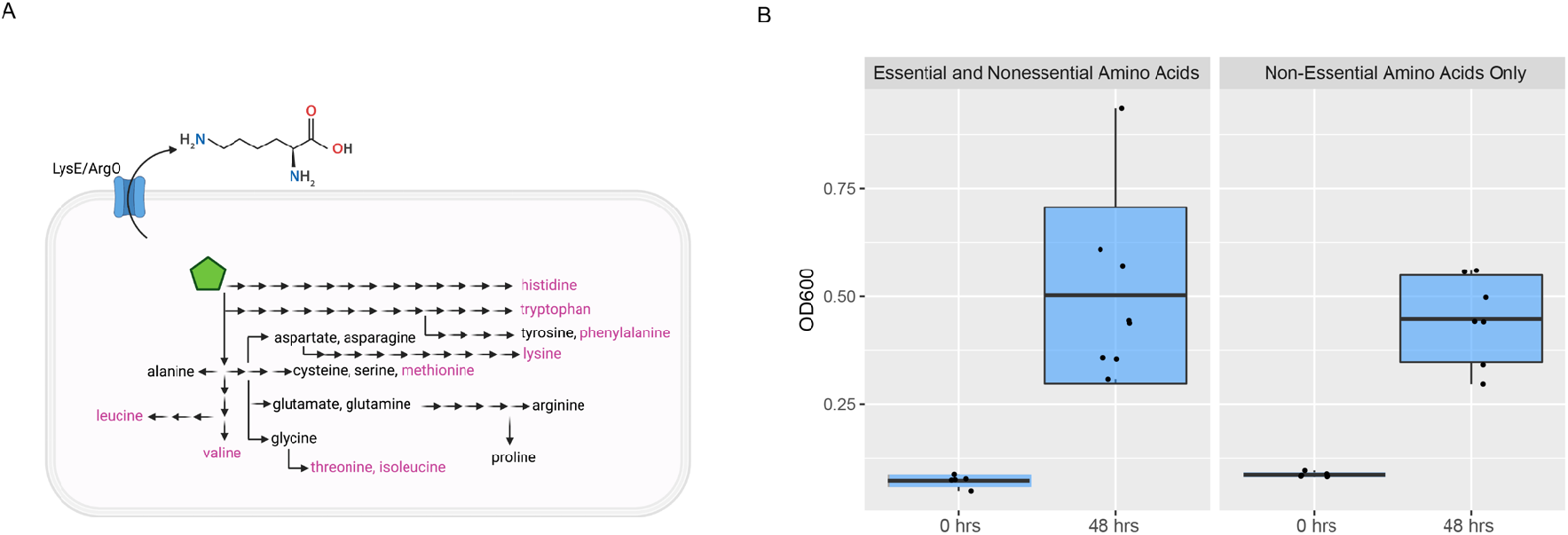
*B. apis* A29 can produce all essential amino acids. **A.** Schematic diagram depicting the amino acid metabolic potential of *B. apis* A29, plus a putative lysine/arginine exporter. Arrows represent enzymatic steps in biosynthetic pathways. Each amino acid that can be synthesized by *B. apis* A29 is positioned at the end of a pathway, with essential amino acids labeled in pink. **B.** Boxplots representing the optical density achieved by *B. apis* A29 after incubating for 48 hours in media containing either all 20 amino acids required for protein synthesis, or only non-essential amino acids. Each group contained at least 5 biological replicates.

To validate the genomic prediction that *B. apis* A29 can synthesize all essential amino acids, we created two minimal media (see methods) containing either all amino acids, or nonessential amino acids only. When provided only nonessential amino acids, *B. apis* A29 showed no growth defects after 48 hours, arriving at a final OD600 similar to that observed when provided all essential amino acids in minimal media, confirming that *B. apis* A29 can synthesize all essential amino acids from nonessential precursors (Figure 2B).

Nutritional symbiosis is most studied in the context of endosymbionts which are often ancient and obligate, resulting in loss of bacterial genes essential for life outside the host (26, 61–64). These losses often result in metabolic dependencies between host and symbiont, or multiple symbionts (11, 12, 65–69). However, in the case of *B. apis,* its position in the larval diet niche appears to maintain selective pressure on free living traits such as complete biosynthetic pathways for the generation of all amino acids (Figure 2A). Since *B. apis* is associated with multiple in-hive environments such as the nurse crop, queen gut, and nectar, it is likely that *B. apis* is faced with a variety of nutritional environments that necessitate metabolic autonomy.

### *B. apis* secretes lysine in the larval diet

After validating our metabolic predictions in culture, we next assessed how the presence of *B. apis* may modify the nutritional composition of the larval diet itself. We reasoned that if *B. apis* were to be supplementing the host with amino acids, it should encode amino acid transporters. Indeed, the *B. apis* A29 genome encodes a LysE/ArgO cationic amino acid transporter (Figure 2A). We therefore used a gene gain/loss analysis across the phylogeny of all sequenced *B. apis* strains and related microbes to identify branches at which amino acid transporters may have been acquired (Figure 3). In the process of performing this analysis, using a larger number of strains, we recapitulated prior results, identifying the acquisition of gluconolactonase, of CRISPR-Cas cassettes, and of several restriction modification systems by *Bombella* (Supplementary Table 2) (56). As we expected, we also identified three orthologs, each encoding putative cationic amino acid transporters, present in *Bombella apis* genomes. The first was present in the progenitor to all these acetic acid microbes (LysE/ArgO). The second was acquired by the ancestor to *Saccharibacter* and *Bombella.* And the third acquired by the ancestor of the *Bombella* clade (ASO19_03265, WP_154981532 and WP_086431440 in the *B. apis* A29 genome LMYH01000007.1; Supplementary Table 3). This result suggests that in the evolution of *Bombella* in honey bee association, the transport of cationic amino acids, such as lysine, was an important trait.

**Figure 3.**
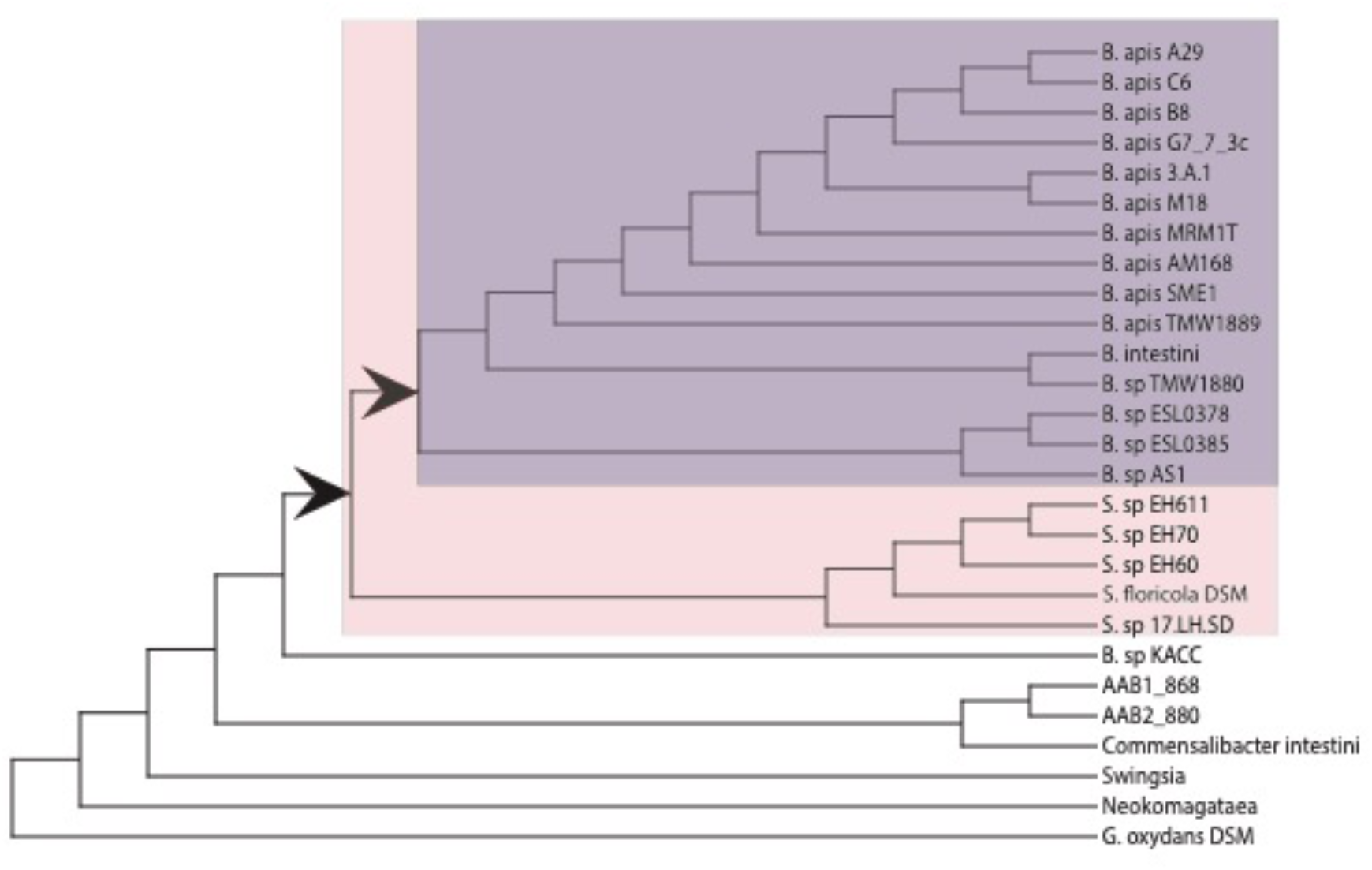
*Bombella apis* strains encode three amino acid transporters, having acquired two cationic transporters in their evolutionary history. Phylogenetic tree generated from conserved core orthologs across the included strains (accessions found in Supplementary Table 1). Predicted acquisitions of cationic amino acid transporters indicated at arrowheads.

We next aimed to experimentally confirm the secretion of lysine by *B. apis* in larval diet. Due to the logistical challenge of acquiring large quantities of natural larval diet, we relied on the synthetic larval diet used in the process of *in vitro* rearing of honey bee larvae, developed by Schmehl et al, 2011. This synthetic larval diet is compositionally like natural larval diet and is sufficient to rear honey bee larvae to adulthood (55). To determine how *B. apis* A29 modifies the larval diet, we, performed gas chromatography-mass spectrometry (GC-MS) on samples of diet incubated with either live or heat-killed *B. apis* A29. We selected this strain because its genome has been previously sequenced, and it grows reliably under laboratory conditions. The heat-killed control allowed us to subtract the nutritional contribution of raw bacterial biomass provided by lysed cells and focus on the output of *B. apis* A29’s active metabolism in the larval diet. Strikingly, larval diet supplemented with live *B. apis* A29 contains significantly higher levels of total essential amino acids (Figure 4A, one-way ANOVA, Tukey HSD, p=0.0006858). Conversely, nonessential amino acids are significantly lower when live *B. apis* A29 is present (Figure 4A, one-way ANOVA, Tukey HSD, p= 0.0298679), while total TCA cycle intermediates are not significantly affected (Supplementary Figure 2, one-way ANOVA, Tukey HSD, p=0.6313555). Though not statistically significant, the TCA cycle intermediate citrate was increased by the presence of live *B. apis* (Supplemental Figure 3, Mann-Whitney U test, Bonferroni correction, p= 0.115346). These patterns suggest an upcycling by *B. apis* of nonessential dietary amino acids into essential amino acids, which are then taken up by the host. The significant increase in total essential amino acids is driven largely by a more than twofold increase in lysine (Figure 4B, oneway ANOVA, Tukey HSD, p<0.0000001). Essential amino acids significantly decreased by live *B. apis* include methionine and threonine (one-way ANOVA, Tukey HSD, p< 0.000001 and p= 0.0006436, respectively)), both of which rely on the same metabolic precursors as lysine. Nonessential amino acids significantly decreased by *B. apis* were glutamate and serine (Supplementary Figure 1, one-way ANOVA, Tukey HSD, p=0.001591 and p=0.0110176, respectively). Though not statistically significant, proline was the only nonessential amino acid increased by the presence of live *B. apis* (Supplemental Figure 1, one-way ANOVA, Tukey HSD, p=0.0866928). Together these data reveal that *B. apis* can dramatically impact the amino acid content of the honey bee larval diet, largely by increasing dietary lysine.

**Figure 4.**
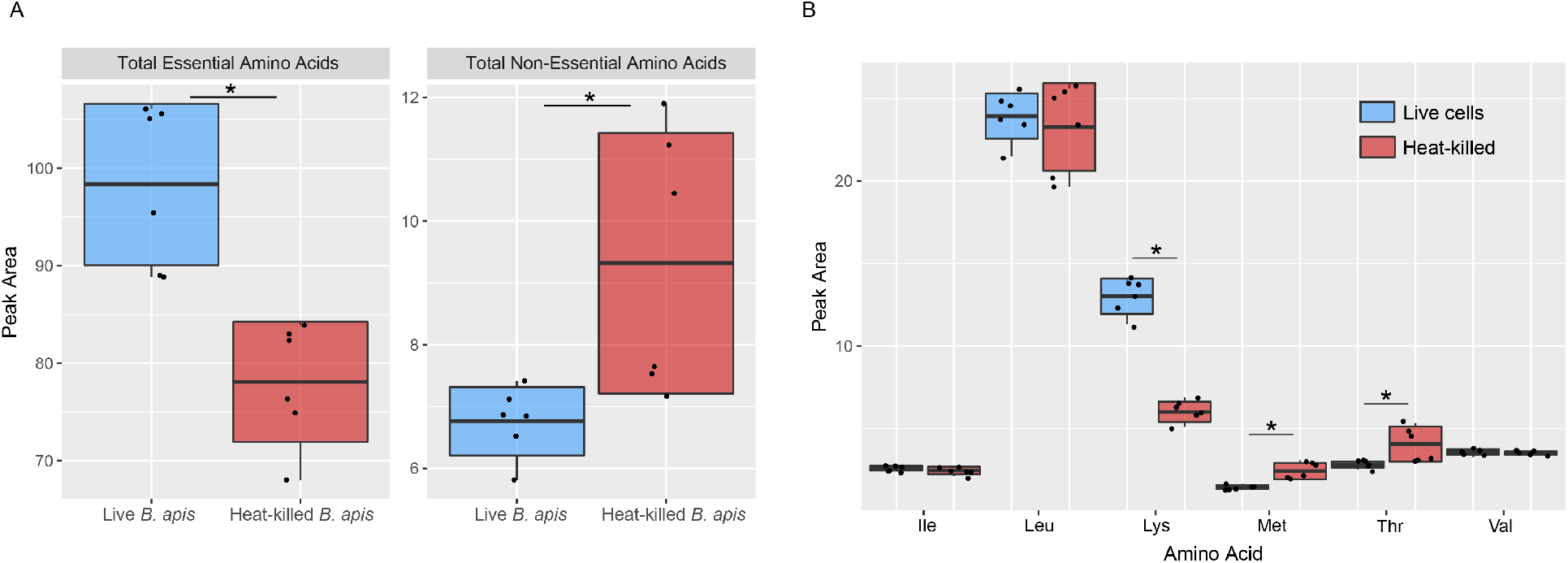
*B. apis* A29 increases the essential amino acid lysine in the honey bee larval diet. **A.** Boxplots showing the total peak areas of essential and non-essential amino acids in synthetic larval diet after incubating with either live (blue) or heat-killed (red) *B. apis* A29. Larval diet incubated with live *B. apis* A29 contained significantly higher total essential amino acids (p=0.0006858) and significantly lower total non-essential amino acids (p=0.0298679). **B.** Boxplots showing the peak areas of individual essential amino acids in synthetic larval diet after incubating with either live (blue) or heat-killed (red) *B. apis* A29. Live *B. apis* A29 results in significantly higher dietary lysine (p<0.0000001) and significantly lower methionine (p< 0.000001) and threonine (p= 0.0006436). Significant differences in peak area were determined using. one-way ANOVA and corrected using Tukey HSD. Prior to ANOVA, data was normalized using Box Cox transformation.

The ability of *B. apis* to not only survive, but meaningfully modify the honey bee larval diet strongly implicates this bacterium as a nutritional mutualist of honey bee larvae. Insects who feed on nutritionally challenging diets must often rely on bacterial partners to supplement missing nutrients (10–13). Many of the best documented examples of bacterial supplementation involve insects gaining essential amino acids from intracellular symbionts (10, 11, 39, 65, 70). *B. apis* A29 appears to be shunting its metabolic energies into production of the essential amino acid lysine, which may be particularly valuable to developing larvae. Lysine appears to play an important role in other holometabolous symbioses; bacterial lysine synthesis and export is crucial for whitefly reproduction, and lysine synthesis is maintained in two *Campotonus* ant symbionts despite genome-wide erosion of central metabolic genes (71, 72). An essential role of lysine on adult honey bee mass was revealed by de Groot in a 1952 study where the author measured adult mass after withholding individual amnio acids from the adult diet. Adult bees deprived of lysine suffered greatly reduced mass relative to those on complete diets (36). Further, many commercial crops which rely on honey bee pollination services only barely meet an adult bee’s minimum lysine requirements (35). Importantly, both studies focused on adult bees, yet we know that the nutritional demands of larvae are more dire (6–8). It is easy, therefore, to imagine a scenario where a honey bee colony must rely on mutualistic bacteria such as *B. apis* to fill in the nutritional gaps in the larval diet.

### *B. apis* bolsters honey bee larval growth under nutrient scarcity

To assess whether the observed metabolic modifications of the larval diet by *B. apis* translate to ecologically meaningful outcomes for honey bee larvae, we conducted an *in vitro* rearing experiment testing the impact of dietary *B. apis* under different diet conditions. Larvae were grafted at 1^st^ instar from naturally mated colonies into sterile multiwell plates containing axenic *in vitro* rearing diet. This approach allowed us to modify the microbial content of the diet as well as the nutrition the larvae received; however we are unable to sterilize field-collected larvae to create axenic individuals. We raised larvae under sterile conditions on either synthetic diet (nutrient-rich) or diet that had been diluted with water by 25% (nutrient-poor) and supplemented them daily with either live *B. apis* A29 or sterile PBS alone, creating four different treatment conditions. Larvae were individually weighed at the end of their larval period, just before pupation and after evacuation of the larval gut. In this experiment, larvae reached masses between 50.9 mg and 158.5 mg (mean 133.4 mg, standard deviation 22.4 mg), well within range of published masses for honey bee 5^th^ instar larvae (73, 74). As expected, dropping the nutritional content of the larval diet by 25% resulted in an average weight drop of 17% in PBS-supplemented control larvae. Therefore, our nutrient limitation treatment did translate to phenotypic differences in the 5th instar larvae. Larvae absent their symbiont but subjected to nutrient-poor conditions were significantly smaller than those in nutrient-rich conditions (Figure 5, Mann-Whitney U test, Bonferroni correction, p=0.013086). In contrast, larvae in the nutrient-poor condition supplemented with *B. apis* A29 were able to reach the same masses as those in nutrient-rich conditions (Figure 5, Mann-Whitney U test, Bonferroni correction, p=0.179). We also noticed that the variance in the masses reached by larvae in the nutrient-poor condition absent their symbiont was significantly greater than those in the nutrient poor condition but given *B. apis* (Figure 5, Levene’s Test of Equal Variance, Bonferroni correction, p=0.017442) – from 53.71 in the *B. apis-*supplemented group to 1000.25 in the un-supplemented group. PBS control larvae in nutrient-limited conditions reached masses as low as 50.9 mg, while those supplemented with *B. apis* under the same nutrient-limited conditions reached more than twice the mass (114 mg for the smallest individual). While there was no statistically significant difference between PBS and *B. apis*-supplemented larvae in the nutrient poor condition (Figure 5, Mann-Whitney U test, Bonferroni correction, p=1), on average, nutrient-poor PBS control larvae were 7% smaller than those supplemented with *B. apis.* Overall, these results indicate that *B. apis* can rescue growth under nutrient limitation. Though nutrient-limited larvae in the PBS group were smaller at prepupation, developmental time was the same between all groups, as indicated by the purging of gut contents on the fifth day of feeding. Coupled with our metabolomic findings, these data showing a growth buffering phenotype of larval bees experiencing poor nutrition indicate a role for *B. apis* as a nutritional mutualist of honey bee larvae.

**Figure 5.**
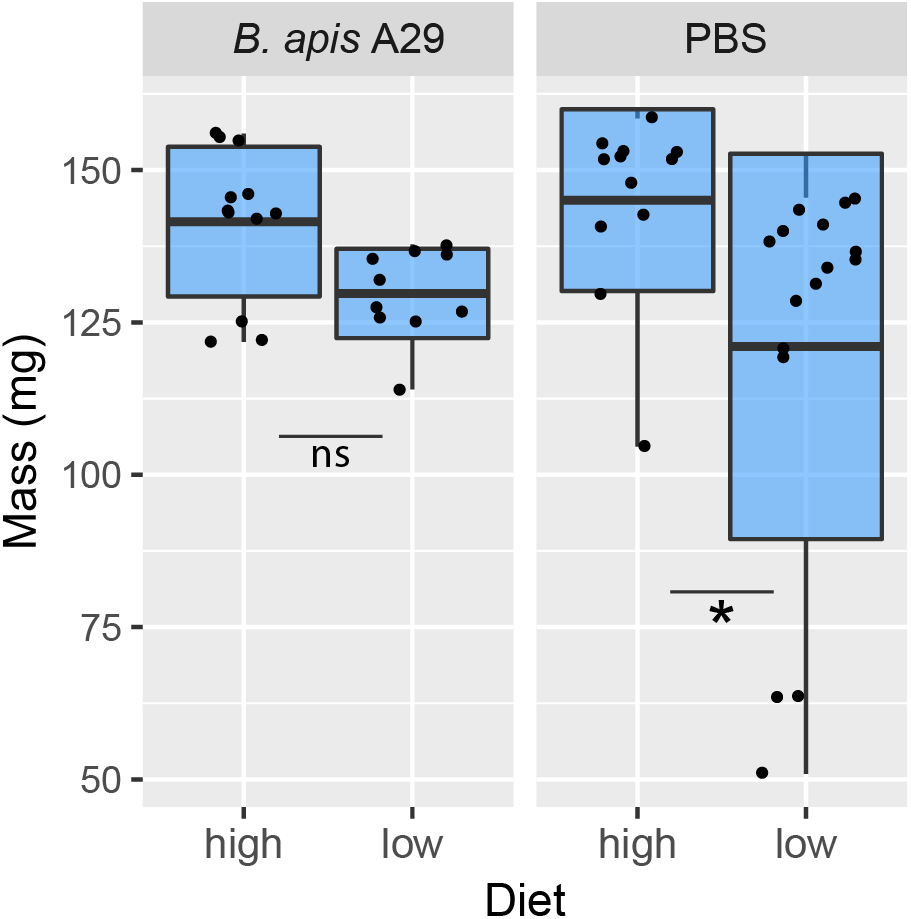
*B. apis* A29 buffers larval mass against poor diet. Boxplot showing the masses of individual larvae after receiving either synthetic larval diet or diet diluted 25% with water, plus either live *B. apis* A29 or sterile phosphate buffered saline (PBS). Larvae given diet supplemented daily with *B. apis* A29 show no significant difference in mass between full or diluted diet (p= 0.19266). Among larvae given PBS only, those receiving diluted diet are significantly smaller than those receiving undiluted diet (p= 0.013086). Significant differences in mass were determined using the Mann-Whitney U test with Bonferroni correction.

### Conclusions and future work

*Bombella apis* is the only honey bee larvae associated microbe that can survive in royal jelly. It synthesizes all essential amino acids and secretes lysine in larval diet. The presence of *B. apis* bolsters honey bee larval mass during nutrient scarcity, which can have dramatic downstream consequences for honey bee colony health. All these data point to the importance of *B. apis* in a colony. Importantly, however, we have not linked the lysine secretion directly to honey bee nutritional supplementation. Future work will focus on genetic modification of *B. apis* to squarely implicate the cationic amino acid transporters and/or amino acid biosynthetic pathways to honey bee nutrition. Also, although we have performed a comparative genomic analysis on multiple strains and determined that all strains are capable of synthesizing and secreting lysine, it remains to be determined whether there is variation in their nutritional bolstering. It is conceivable, that some strains are better mutualists than others, and the link between genetic and phenotypic diversity in *B. apis* is an active area of research in the lab. Indeed, based on our data (Figure 1), we might suspect that different strains are better able to survive and supplement larvae in the royal jelly diet. Additionally, genetic differences in the honey bee larvae in our experiments may account for some of the variance we observed in our experiments, although we were careful to randomize our sampling of larvae across treatments. The interaction between honey bee genetics and the microbiome is only starting to be explored and it would be a benefit in future experiments to at least control for genetic variation in a colony by using single drone inseminated queens. Finally, our experiments were performed on *B. apis* alone and it is likely that *Lactobacillus kunkeei* also plays some role in larval development given that 1) it is routinely isolated from larval niches and honey bee hive environments and 2) it can survive in the presence of some royal jelly. Understanding the interaction between *B. apis* and *L. kunkeei* will be important to understanding the role that these microbes play in honey bee larval development and nutritional supplementation.

## Methods

### Culturing bacteria

*Bombella apis* A29 was grown at 34°C, ambient oxygen, shaking at 250 rpm, in Bacto-Schmehl (BS) liquid media. BS growth media is derived from the larval diet outlined in Schmehl et al., 2011 conceived for growth of the honey bee larvae and is described in the following section. The designation BactoSchmehl is used with the written permission of Dr. Daniel Schmehl.

### Bacto-Schmehl (BS) Media

- 5% w/v D-Glucose
- 5% w/v D-Fructose
- 1% w/v Yeast Extract
- 4% v/v 5X Sigma M9 Salt Solution (catalog #M9956)
- 0.2% v/v Cation solution (1% [v/v]; 100 mM MgSO4 and 10 mM CaCl2 in diH_2_O)
- 84.8% Milli-Q H2O

Cation solution must be autoclaved separate from the M9 salt solution. BS media final pH 6.5.

### Minimal Bacto-Schmehl Media (mBS)

- D-Glucose
- D-Fructose
- 5X Sigma M9 Salt Solution (catalog #M9956)
- Cation solution (1% [v/v]; 100 mM MgSO4 and 10 mM CaCl2 in diH_2_O)
- 1X Sigma MEM Vitamin Solution (catalog #M6895)
- 1X Sigma MEM Amino Acids solution (catalog #M5550) or 1X Sigma Non-essential Amino Acid Solution (catalog #M7145)
- Milli-Q H2O

### Bacterial growth in honey bee larval diet

Overnight cultures of each bacterial strain were grown in BS broth *(Bombella apis* A29, *B. apis* B8, *B. apis* C6, *B. apis* SME1, *B. apis* MRM1T, *Lactobacillus kunkeei* AJP1, and *Fructobacillus fructosus* AJP3). Cultures were washed (cells spun down in microcentrifuge tubes and resuspended in sterile PBS) twice in sterile 1X PBS, then normalized to 10^7^ CFU/ml. 50 ul of each bacterial suspension was added to 500 ul of BS media with an increasing proportion of commercial royal jelly up to 50%, in triplicate. After 24 hours incubating at 34°C, samples were serially diluted and plated on BS agar, in triplicate. CFUs were counted to determine numbers of viable cells.

### Comparative genomic content analyses, identification of orthologs, and gain/loss analysis

To define orthologs, protein sequences were extracted from NCBI annotated sequence files for the *Acetobacteraceae* clade rooted on *Gluconobacter* (Figure 3). Reciprocal best blast hits were calculated, and genes clustered into ortholog groups using complete linkage. Conserved core orthologs were used to generate the species tree for these genera and this was used, in conjunction with GLOOME (75) to infer branch-specific gene gain/loss events (Supplementary Table 2). To define presence/absence of amino acid biosynthesis genes (Supplementary Table 1), ortholog representatives were run against GapMind (76) to find amino acid biosynthetic genes in the proteomes. Additionally, annotation based on NCBI’s PGAP was used, in conjunction with DOE’s IMG/M to confirm the putative function of orthologous groups of genes.

### Minimal media assay

An overnight culture of *B. apis* A29 was grown in BS broth, then washed in sterile 1X PBS before inoculating mBS media containing either a complete amino acid solution or a non-essential amino acid solution (see Methods). Cultures were diluted 1:100 in mBS and incubated at 34°C for 48 hours. Optical density (OD600) was measured using a spectrophotometer at the start of the experiment and at 48 hours.

### Larval collection and rearing

All larvae were collected from hives at the IURTP Bayles Road field site in October 2019. To minimize any effects of manipulation of the bee larvae and of genetic differences between colonies, all larvae were evenly distributed between treatments with respect to colony of origin and order of collection.

First instar larvae were grafted from comb using a plastic grafting tool and were deposited into plastic queen cups in 48-well plates. Grafting and rearing protocols were conducted according to the Schmehl et al. 2011 protocol with several deviations noted here. Each larva was grafted into 10 ul UV-sterilized larval Diet A (Schmehl et al., 2011), at the field site over the course of two hours. The larvae were transported back to the laboratory and incubated in darkness at 34°C and 90% RH. After overnight incubation, the larvae that did not survive grafting were removed and the remaining larvae were divided into experimental groups.

All larvae were fed according to the diet recipes of Schmehl et al., 2011. Larval diet was made no more than 48 hours in advance of each feeding. All diet was UV-sterilized for 20 minutes to remove any potential bacterial contamination. Diet was refrigerated at 4°C between feedings and warmed to 34°C prior to each feeding. Larval diet was dispensed to individual larvae under a laminar flow hood using sterile pipettes. Larvae were fed mid-afternoon across all experiments.

The larval feeding and bacterial supplementation timeline is outlined in the table below:

|  |  |
| --- | --- |
| Day 0 | Larvae grafted from field |
| Day 1 | 10 ul Diet A |
| Day 2 | 20 ul Diet B, 5 ul bacterial cell suspension |
| Day 3 | 30 ul Diet C, 5 ul bacterial cell suspension |
| Day 4 | 40 ul Diet C, 5 ul bacterial cell suspension |
| Day 5 | 50 ul Diet C, 5 ul bacterial cell suspension, diet samples collected for metabolomics |
| Day 6 | Prepupal masses recorded |

In all supplementation experiments, high-diet larvae received undiluted larval diet according to Schmehl et al., 2011 recipes. Low-diet larvae received larval diet from the same batches, divided and diluted by 25% using sterile deionized water. Each batch of diet was divided and diluted prior to UV-sterilization.

On the sixth day following larval grafting, larvae had consumed all remaining diet and defecated in their cells. Each larva was then removed from its cell and individually weighed. Residual diet and excrement were removed from the surface of each larva using a modified plastic grafting tool prior to weighing. Any larvae that were accidentally punctured during the weighing process were not weighed.

### Bacterial supplementation of larval diet

An overnight culture of *B. apis* A29 was washed twice to remove excess media, then resuspended in sterile PBS. Optical density (OD) was measured using a spectrophotometer to confirm adherence to known OD/CFU. Prior to larval feeding, bacterial suspensions were normalized to 10^4^ CFU/ml using PBS. For experiments involving heat-killed controls, this normalized solution was then divided, and half was subjected to boiling for 10 minutes. 5 ul of bacterial suspensions or PBS was pipetted into each queen cup containing a single larva. Bacterial suspensions were given immediately after daily feeding. All feeding and supplementation was performed using sterile technique under a laminar flow hood. Larval masses were compared in R using pairwise Mann-Whitney U-tests, then Bonferroni corrected for multiple comparisons

### Metabolomic analysis

On the fifth day following larval grafting, samples were taken from the larval diet of *in vitro* reared larvae for metabolomic analysis. 8 hours after diet administration and bacterial supplementation, 3 ul of diet was removed from each larval cell. Samples were combined based on treatment, yielding 12 ul samples representing diet from four individual larvae. These 12 ul samples were immediately flash-frozen in liquid nitrogen and stored at −80°C before GC-MS. Samples were randomized prior to GC-MS to control for variation between individual GC-MS runs.

GC-MS analysis of larval diet samples were conducted using a modified version of a previously described method (77). Briefly, 12 mL of larval diet was dissolved in 800 mL of prechilled (−20 °C) 90% methanol containing 2 μg/mL succinic-d4 acid. The sample was incubated at −20°C for 1 hour and centrifuged at 20,000 x g for 5 minutes at 4°C. 600 ml of the supernatant was transferred into a new 1.5 mL microcentrifuge tube and dried overnight in a vacuum centrifuge. Dried samples were resuspended in 40 μL of 40 mg/mL methoxylamine hydrochloride (MOX) dissolved in anhydrous pyridine and incubated at 37°C for 1 hour in a thermal mixer shaking at 600 rpm. Samples were then centrifuged for 5 minutes at 20,000 x g and 25 μL of supernatant was transferred into an autosampler vial with a 250 μL deactivated glass microvolume insert (Agilent 5181-8872). 40 μL of N-methyl-N-trimethylsilyltrifluoracetamide (MSTFA) containing 1% TMCS was then added to the sample, at which point the autosampler vial was capped and placed at 37°C for 1 hour with shaking (250 rpm).

1 μL of sample was injected an Agilent GC7890-5977 mass spectrometer equipped with a Gerstel MPS autosampler. Samples were injected with a 10:1 split ratio and an inlet temperature of 300°C. Chromatographic separation was achieved using a 0.25 mm x 30 m Agilent HP-5ms Ultra Insert GC column with a helium carrier gas flow rate of 1.98 mL/min. The GC temperature gradient was as follows: (1) Hold at 95°C for 1 min. (2) Increase temperature to 110°C with a 40°C/min ramp. Hold 2 min. (3) Increase temperature to 250°C with a 25°C/min ramp. (4) Increase temperature to 330°C with a 25°C/min ramp. Hold for 4 minutes. Extraction and GC-MS was performed by the Indiana University Mass Spectrometry Facility. Metabolite concentrations were compared in R using pairwise Mann-Whitney U-tests, then Bonferroni corrected for multiple comparisons, or were normalized using Box Cox transformation prior to one-way ANOVA and Tukey HSD correction.

