## Supplementary Figures and Legends for "A honey bee symbiont buffers larvae against nutritional stress through lysine supplementation"

Supplementary Table 1 – All sequenced *B. apis* strains retain the ability to synthesize all amino acids. Table generated from conserved core orthologs across the included strains showing presence/absence of amnio acid biosynthesis genes. ‘oid’ refers to the ortholog ID in our analysis of orthologous genes, ‘Name’ refers to the amino acid biosynthesis gene annotation, ‘Pathways/steps/scores’ refers to the biosynthetic pathway in which each gene is found, the enzymatic step in the pathway, and the GapMind score. GapMind score is either 2 (high confidence), 1 (medium confidence), or 0 (low confidence). In the columns below each sequenced strain, ‘1’ means that a given gene was identified in the corresponding genome and ‘0’ means that it was not identified.

Supplementary Table 2- All *B. apis* genomes contain multiple cationic amino acid transporter orthologs. Gene gain/loss analysis showing all the gains and losses across the phylogeny of all sequenced *B. apis* strains and related microbes in Figure 3. ‘oid’ refers to the ortholog ID in our analysis of orthologous genes, ‘Name’ refers to proteins identified across all genomes analyzed. In the columns below each sequenced strain, ‘g’ means that a given gene was gained by that strain and ‘l’ means that it was lost.

Supplementary Table 3 – Accession numbers for all sequenced strains used in this work.

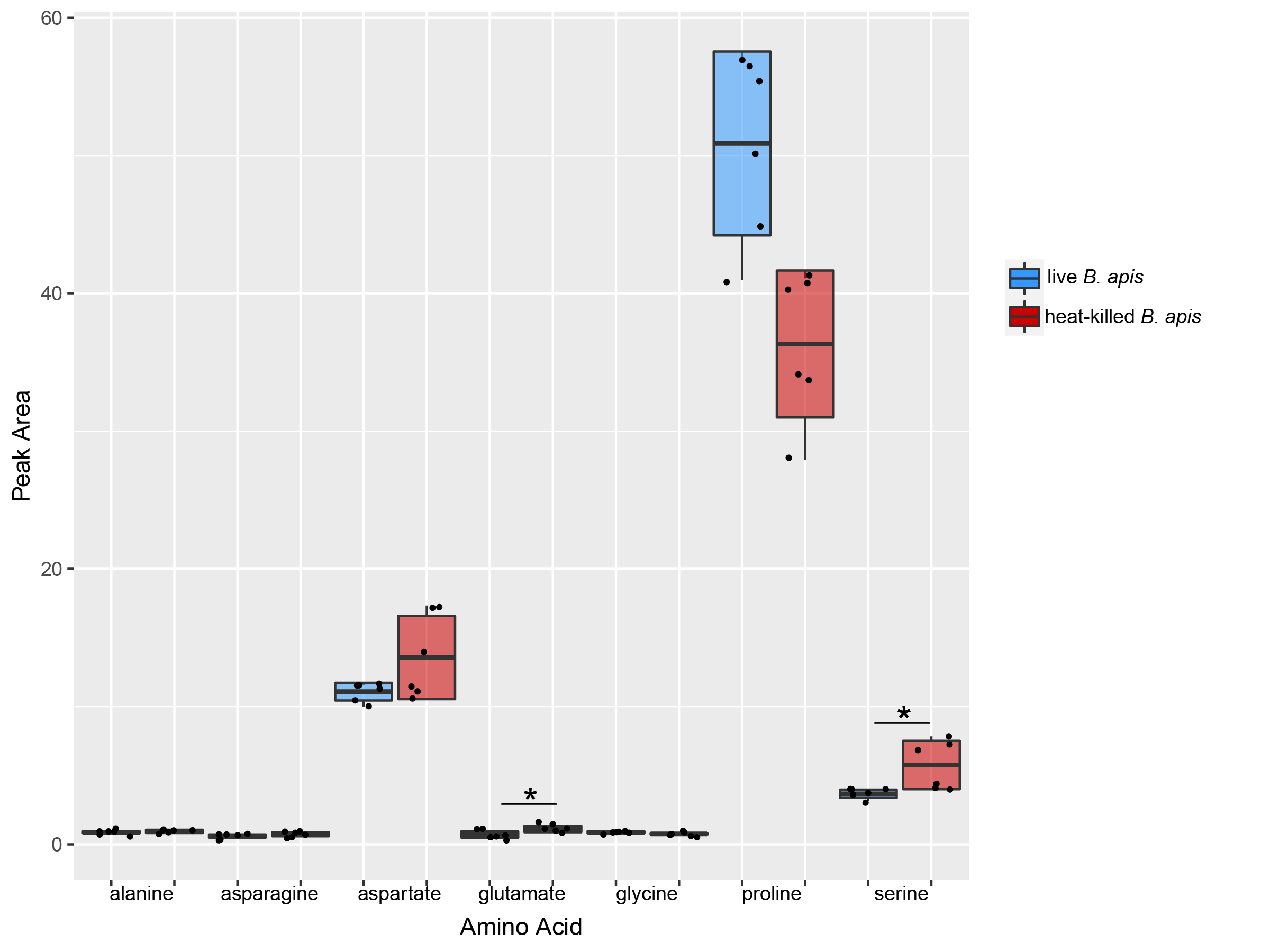

Supplementary Figure 1 – The nonessential amino acid landscape of the honey bee larval diet is modified by *B. apis* A29. Boxplots showing the peak areas of individual nonessential amino acids after incubating with either live (blue) or heat-killed (red) *B. apis* A29. Live *B. apis* A29 results in significantly lower dietary glutamate (p=0.0015910) and significantly lower serine (p=0.0110176). While not significant, *B. apis* increases dietary proline (p=0.0866928). Significant differences in peak area were determined using one-way ANOVA and corrected using Tukey HSD. Prior to ANOVA, data was normalized using Box Cox transformation.

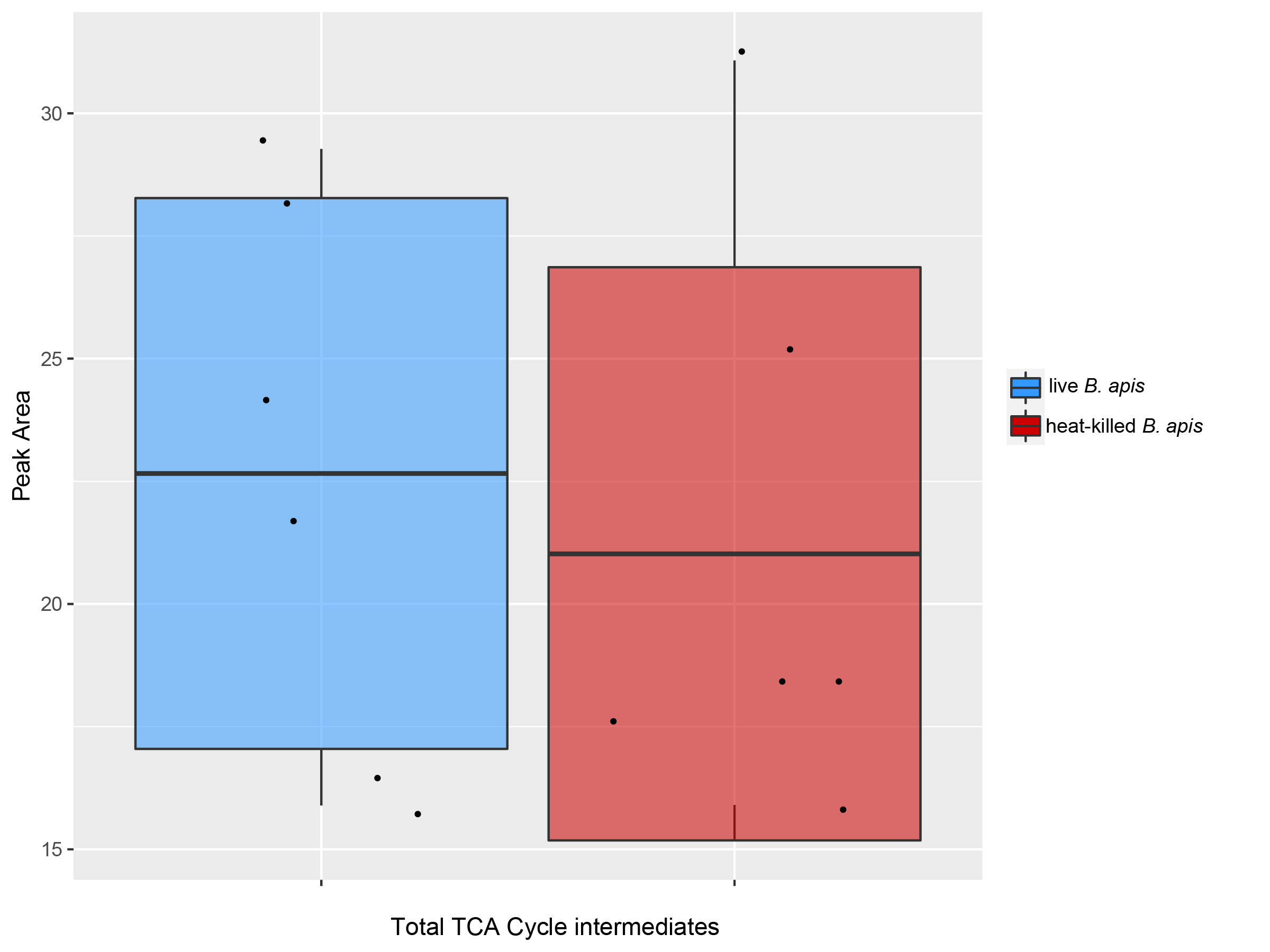

Supplementary Figure 2 - *B. apis* A29 does not significantly affect total dietary TCA cycle intermediates. Boxplot showing the total peak areas of TCA cycle intermediates in synthetic larval diet after incubating with either live (blue) or heat-killed (red) *B. apis* A29. Significant differences in peak area were determined using one-way ANOVA and corrected using Tukey HSD.
